# An automatic and efficient pulmonary nodule detection system based on multi-model ensemble

**DOI:** 10.1101/2020.04.14.040931

**Authors:** Jian Chen, Wenlei Wang, Bin Ju, Jing Jiang, Liying Zhang, Huilin He, Xuanxuan Zhang, Yuqiang Shen

**Affiliations:** Zhejiang University School of Medicine Fourth Affiliated Hospital, Yiwu, 322000, China; The Wowjoy Technology, Hangzhou, 310006, China

**Author notes:** These authors contributed equally to this work (Jian Chen, Wenlei Wang, Bin Ju).

## Abstract

Accurate pulmonary nodule detection plays an important role in early screening of lung cancer. Although there are many presented CAD systems based on deep learning for pulmonary nodule detection, these methods still have some problems in clinical use. The improvement of false negatives rate of tiny nodules, the reduction of false alarms and the optimization of time consumption are some of them that need to be solved as soon as possible. In view of the above problems, in this paper, we first propose a novel full convolution segmentation framework for lung cavity extraction in preprocessing stage to solve the time consumption problem of the existing pulmonary nodule detection systems. Furthermore, a 2D-NestedUNet segmentation network and a 3D-RPN detection network is stacked to get the high recall and low false positive rate on nodule candidate extraction, especially the recall of tiny nodules. Finally, a false positive reduction method based on multi-model ensemble is proposed for the further classification of nodule candidates. Our methods are evaluated on several public datasets, LUNA16, LNDb and ChestCT2019, which demonstrated the superior performance of our CAD system.

## Introduction

Lung cancer is a leading cause of cancer-related death both in men and women. Every year lung cancer results in about 170 million deaths worldwide. The early diagnosis of a lung lesion is recognized as the most important method to reduce the lung cancer mortality rate. The low-dose CT scans which is one diagnostic way can be used to screen for lung cancer in people. Using this screening method can decrease the risk of dying from lung cancer. Now researchers are looking for new attempts to refine CT screening to better predict whether cancer is present [1]. Due to the rich pulmonary vascular structure and the different skill levels of radiologists, the potential malignant lesions are easy to be ignored. In the clinic, an effective way to deal with this problem is to diagnose by two radiologists respectively and then to summarize their answers. However, this may increase the workload of radiologists [2]. With the development of digital medical image processing, using computer-aided diagnosis (CAD) system to assist pulmonologists in clinical diagnosis is a trend. However, the traditional approaches are less accurate and more complicated in detecting pulmonary nodules at early stage of lung cancer so that are hardly applied to help doctors in clinical diagnosis.

In recent years, with the rapid development of deep learning in the field of medical image analysis, a large number of CNN-based CAD systems have been utilized for the detection of pulmonary nodules [3–5]. Compared with the traditional methods [6, 7], the DNN-based methods have made great progress in accuracy and practicability of pulmonary nodule detection. Nevertheless, many challenges still exist in the lung nodule detection procedure. The detection procedure is usually divided in three stages, i.e. image prepossessing, nodule candidates extraction and false positives reduction. Firstly, in prepossessing step, how to quickly and efficiently extract lung cavity plays an essential role. Due to the various sizes and morphology of pulmonary cavity, the traditional multi-threshold-values based segmentation is not robust enough. Therefore, offering an efficient lung cavity extraction method is very important. Secondly, in clinical practice, radiologists pay more attention to small nodules, because these small nodules are more likely to cause lung cancer in the future. To ensure high sensitivity for them, in other words, CAD is needed to have a high recall rate for small nodules that may better resolve lung cancer early screening. However, the size of small nodules is too small and their radiographic manifestations always appear ground glass attenuation so that it is likely to be neglected. The presented methods, neither using 2D nor 3D system, can find small pulmonary nodules well [3, 4]. That means the developers need invest more research in this area in the future. Thirdly, because abundant tissues exist in the human pulmonary cavity, for instance blood vessel and chest wall, the appearances of these tissues are very similar with the appearance of the pulmonary nodules. This results in producing amounts of false alarms during detection. Making something to accurately distinguish between tissues and nodules thereby reducing the number of false positive signals is very important and necessary [8].

The primary aim of this study is to develop an advanced CAD system that extracts information from medical images efficiently and provides radiologists a precise and timely diagnosis of lung lesion. The key contributions of this paper are summarized below:

- we present a FCNN framework which is based on U-Net [12] for quickly and stably performing pulmonary cavity extraction.
- we stack a 3D-RPN based detection network [3] and a 2D-NestedUNet based segmentation network [13] for providing target candidates to satisfy high recall rate and low false positive rate of pulmonary nodule detection.
- we propose an integrated classification network which consists of ResNet [23], DenseNet [24] and SENet [25], summarizing the results from detection step and classification step to reduce false candidates.

## Related Work

Before the rapid development of CNN in medical image processing, the pulmonary nodules detection was mostly based on hardcrafted features. In recent years, with the successful application of deep learning in medical images, the intelligent screening system of pulmonary nodules has been greatly developed. Based on the exploited deep architecture, these approaches can be divided into proprecessing, candidate nodules extraction and false positive reduction. In [3, 4, 14], the pulmonary cavity of the CT scans was obtained by traditional image processing methods such as region growth and morphological operations. [15] proposed the spot detection for extracting the nodule candidates. Using this method, nodule and non-nodule samples could be distinguished according to their different shape, size, specific texture, density and other features. However, due to the choices of thresholds, the detection performance of this method for small nodules and tiny nodules was very poor and not robust. In recent years, many excellent CNN-based CAD systems for automatic pulmonary nodules detection are presented, which can roughly be divided into 2D image slices and 3D volume images. In [3], 2D image slices were fed into Faster-RCNN [16] to locate suspected nodules and output their sizes in CT images. First from the perspective of 3D volume images, [4] established a pulmonary nodules detection system with the aid of 3D-Faster-RCNN framework. This method could make full use of the spatial information of 3D images. In [17], the attention mechanism is brought into the approach presented in [4] to further acquire 3D spatial semantic information. In [18], a new self-supervised pulmonary nodule detection framework is proposed based on 3D-RPN to improve the sensitivity of nodule detection by adopting multi-scale features to increase the resolution of nodule cubes. Compared with the 2D models, these 3D methods employ spatial semantic information to better analyze the morphological characteristics of nodules and locate them much accurately. However, their recalls for small nodules are not high so that would be easy to increase the false negatives rate for small nodules and nodules. [19] presented a 3D full convolutional network which is based on V-Net [20] to extract nodule candidates. A high recall is the advantage of this solution. For reducing more false positives, [4, 6] proposed a simple 3D convolutional network adding the back of detection module to filter false candidates. [21] introduced the attention mechanism after false positive reduction module to facilitae the performance of classification for pulmonary nodules (benign and malignant).

## Methods

Our proposed CAD system can be roughly divided into three stages: 1) FCNN based lung segmentation, 2) multi-model ensemble based pulmonary nodule extraction, and 3) false positive reduction (the whole pipeline shown in Fig 1).

**Fig 1.**
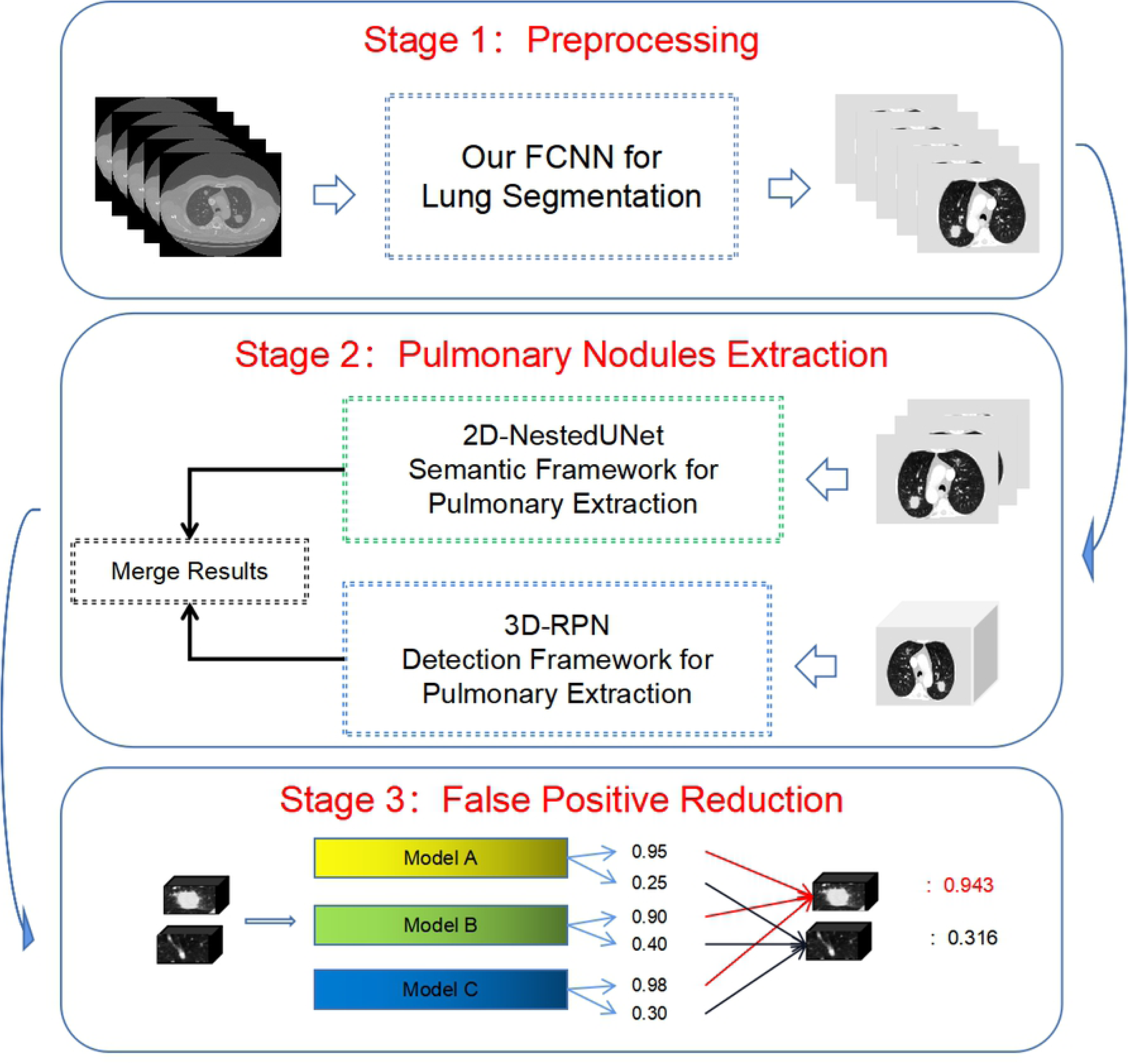
The Overview of Our CAD system. A whole CT scan is fed into our system to predict nodule candidates. The process consists of three stages, i.e. preprocessing, pulmonary nodules extraction and false positive reduction. In first stage, raw data will be processed to extract lung cavity and then fed into second stage to find pulmonary nodule candidates. Finally, the candidates will be filtered in stage 3 to reduce false alarms.

### Novel FCNN for Lung Segmentation

Accurate lung segmentation is the basics for rapidly constructing automated pulmonary nodule detection system with high-accuracy and sensitivity. By using the traditional method, the 2D single slice is first processed with a Gaussian filter to remove the fat, water and kidney background and then followed with a 3D connection region extraction module to remove irrelevant regions. However, this method is time consuming and unstable. In order to accelerate this processing for experiment and deployment, this paper propose a lung segmentation network, which is based on U-Net. The encoder path includes five consecutive convolution modules, the number of feature maps is doubled at each convolution module. Each convolution module consists of two convolution filters with kernel size = 3, followed by a max pooling function with kernel size = 2 and stride = 1. Each convolution filter is followed with a BatchNormalization and a nonlinear activation function ReLU. Finally, the final layer in the encoder path produces a high dimensional image representation with high semantic information.

Assume that the input of the segmentation network is a single-channel grayscale image with a size of 1×H×W, denoted as *I*_*in*_, and the output of each convolution module *i*^*th*^ in the encoder path is denoted as 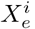, which represents the number of output channels, each convolution module *i*^*th*^ takes as input to 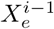. At the end of the encoder, a feature map *F*_*e*_ will be obtained whose size is reduced by 4 times compared to *I*_*in*_.

Different from the traditional U-Net structure, we make some modification for our FCNN in the convolution mode of the decoder path, which is to use the same convolution structure as the encoder path to implement feature decoding, and at the end of the decoder, a convolution filter with kernel size = 1 × 1 used as the network output layer. The entire network will eventually output a segmentation prediction result with the same size as the input image. The predicted value of each pixel represents the likelihood that this pixel belongs to a lung cavity. The overall lung cavity segmentation network is shown in Fig 2, using the Dice coefficient as loss function which is same to original U-Net [12].

**Fig 2.**
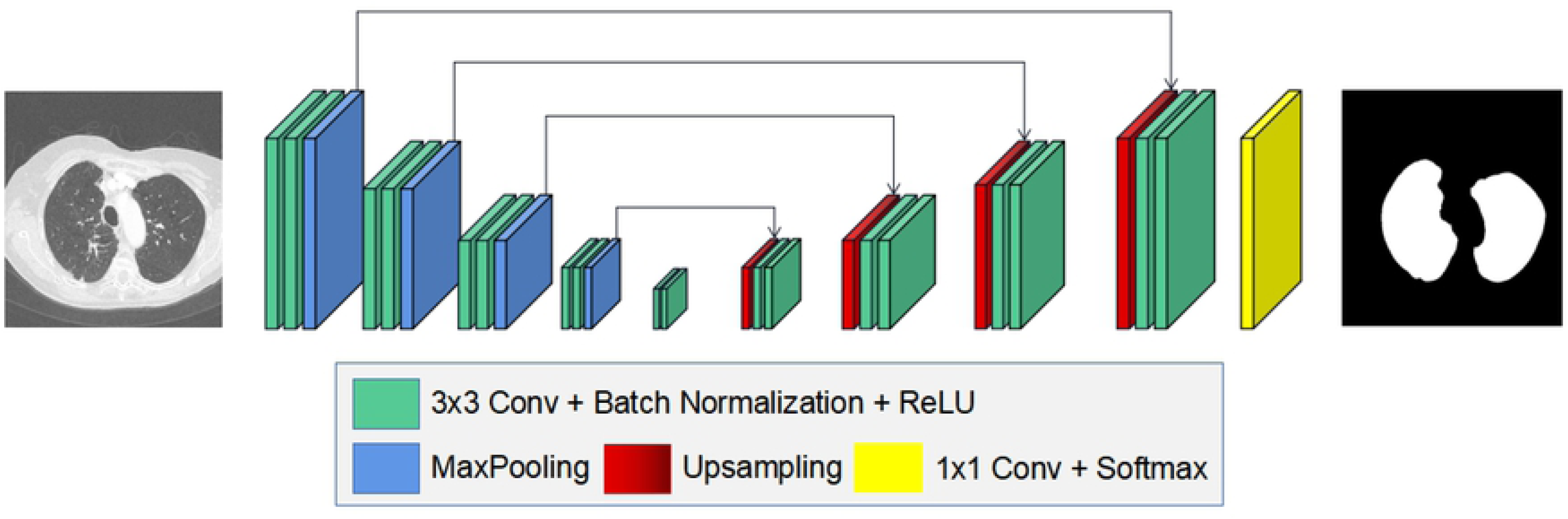
The Schematic of Our FCNN framework for Lung Segmentation.

### Nodule Candidate Extraction based on Multi-model Ensemble

In this section, we design a multi-model ensemble network, which is combined with a 2D segmentation network and a 3D detection network to sensitively screen the nodule candidates.

#### 2D NestedUNet for Pulmonary Nodule Extraction

NestedUNet [22] can accelerate network optimization by using dense convolutional blocks which bridge the semantic gap between the feature maps of the encoder and decoder. However, we found that NestedUNet with depth supervision module is difficult to be optimized and easy to cause gradient explosion while training for pulmonary cavity segmentation in this work. Thus we use the fast version of NestedUNet with the same loss function from the original NestedUNet as our segmentation module, the specific formulas are summarized bellow:

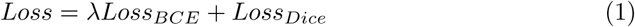

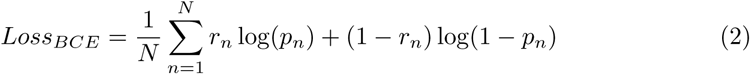

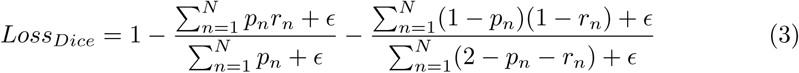

where *p*_*n*_ is the probability that pixel n is predicted as lung cavity, *r*_*n*_ is the true category of pixel n, *r*_*n*_ = 1 for the lung cavity and, *r*_*n*_ = 0 is regarded as backgound. *n* ∈ {1, 2, 3 … *N*}, *N* = *H* × *W*, *ϵ* is a minimal number.

#### 3D RPN for Nodule Extraction

Pulmonary nodule detection, as a crucial step in the CAD systems, aims to accurately locate the nodule, meanwhile catching more true nodule candidates and reducing the number of non-nodule candidates as much as possible. We present a 3D RPN model for detecting nodule candidates from CT images, where a modified U-Net is the backbone network [4]. The structure of pulmonary nodule detection is shown in Fig 3. The network consists of an encoder and a decoder. Due to the limitation of GPU memory, the input of the 3D RPN network is cropped from the 3D CT image. The size of the input image is 128×128×128. The encoder includes 3D convolutional layers, 3D residual blocks, and 3D maximum pooling layers. Each 3D residual block contains three residual units. The decoder has deconvolutional layers, combining units and residual blocks. Each combining unit concatenates the output of the convolutional layer and the corresponding deconvolutional layer, which likes the long connection in U-Net. In the left combining unit, the location information is introduced as an extra input, the feature map of this combining unit has the size of 32×32×32×131. Finally, the output feature is resized to 32×32×32×3×5 after two convolutional layers. The last two dimensions correspond to the anchors and regression box respectively. There are three anchors with different scales, corresponding to three bounding boxes with the length of 3, 10 and 20 mm. The loss function is defined as below:

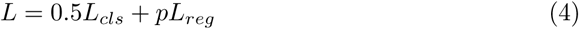

where *p* ∈ {0, 1} (0 for negative and 1 for positive). The classification loss *L*_*cls*_ is the cross-entropy loss and the regression loss *L*_*reg*_ is the smooth L1 loss function.

**Fig 3.**
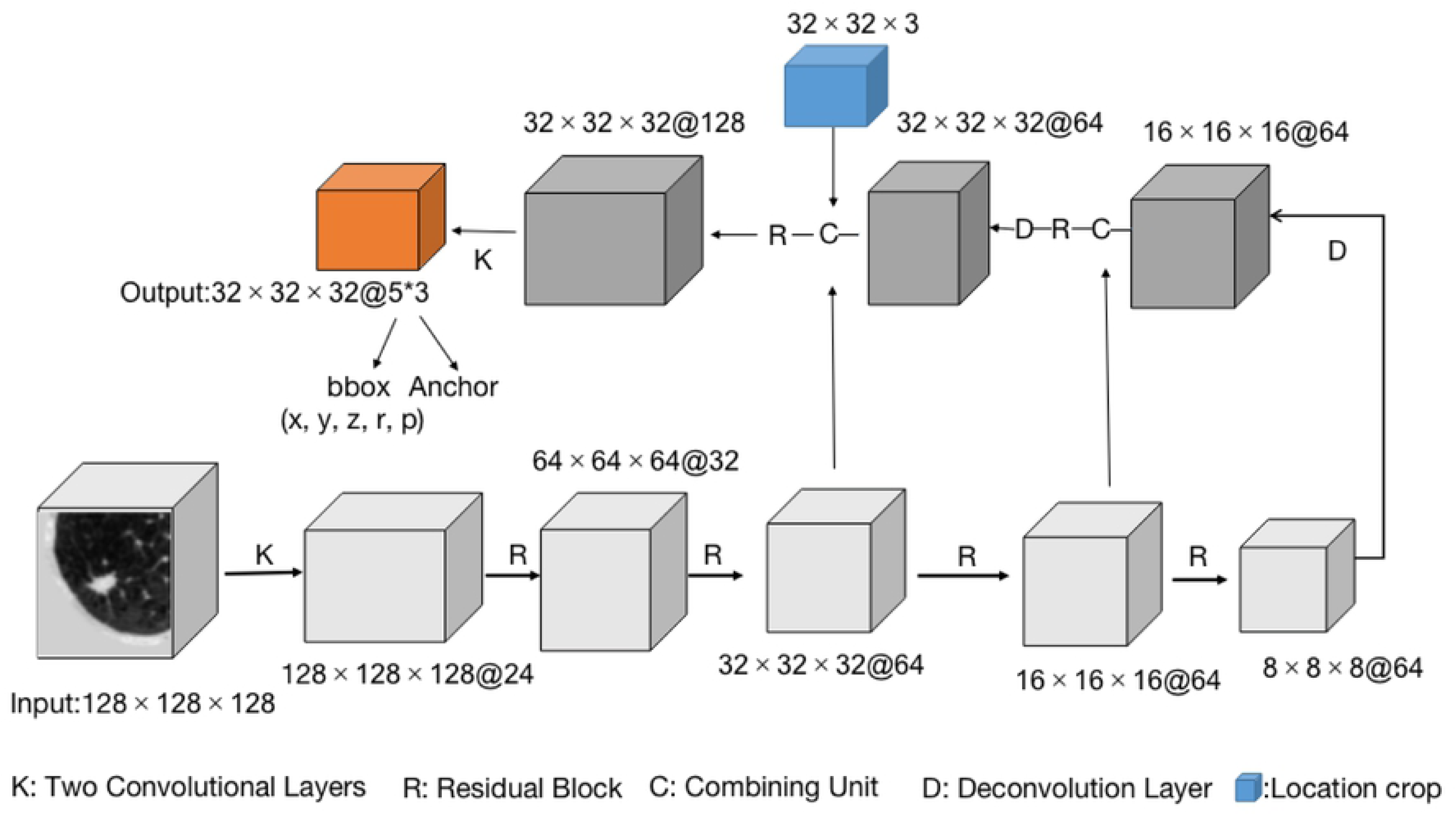
The Structure of 3D RPN for Nodule Detection. Each cube represents a 4D tensor. The size is *Length × Width × Height × Channels*. The last two dimensions of the output tensor correspond to the three anchors and five regression box indicators (nodule coordinates x, y, z, nodule size r and probability).

#### Combination of Nodule Candidates

After a CT scan volume is fed to the aforesaid 2D and 3D pulmonary nodule extraction methods respectively, two lists of pulmonary nodule candidates will be provided, here denoted as *L*_1_ and *L*_2_. Then we can combine these candidates by using algorithm 1. Firstly, based on NMS algorithm, we remove the parts of candidates whose detection boxes have significant overlaps with others. After that, we delete the parts of candidates from 2D segmentation that have high overlaps with above 3D handled results also based on NMS. The 3D detection method has high accurate location as benefit point however its recall rate is relatively lower than 2D segmentation method. Meanwhile, using 2D segmentation network will provide a large number of false positive candidates. Combing the results of two different methods can help us to get accurate position of pulmonary nodules and avoid excessive false alarms. That is the reason why we use model ensemble strategy. Besides, based on CUDA platform, we use multi-thread technology to fast perform the calculation during the comparison process so that can dramatically reduce the time cost of fusion phase.

### False Positives Reduction

As previously mentioned, nodule candidates from detection phase may contain many false alarms, although we have taken some measures to suppress this issue. To reduce the non-nodule candidates, we propose a 3D DCNN-based false positive reduction module which contains three network models. Fig 4 shows our architectural diagram for false positives reduction, the module is based on the 3D network of ResNet [23], DenseNet [24] and SENet [25]. For the center of a given nodule candidate, we extract a 3D data cube of size 32 × 32 × 32 that possibly includes pulmonary nodule as input to the three network models, and then get the probabilities *p*_*i*_ (i ∈ {1, 2, 3}) of the three classifiers. To further eliminate false positives, we design the strategy described as

#### Algorithm 1 Merging two nodules results extraction

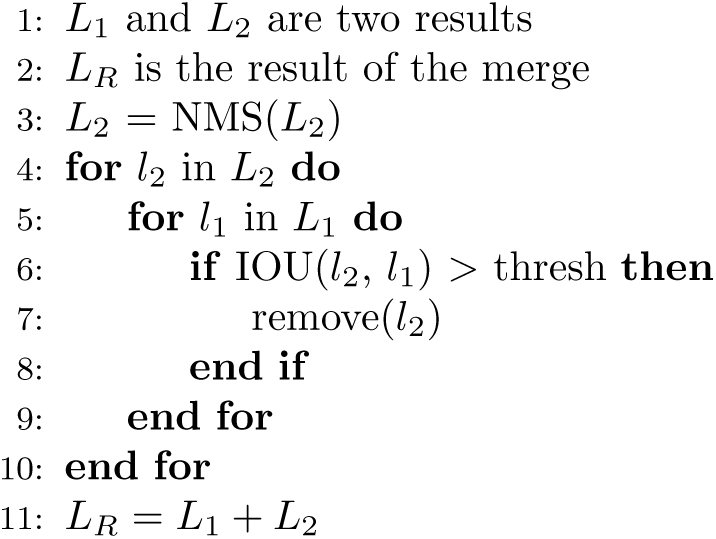

follows: 1) for predicted probability *p*_*i*_, if *p*_*i*_ ≥ *T*_1_, the final classify probability of the input cube is equal to the average value of all *p*_*i*_ ≥ *T*_1_; otherwise, the final result is the average value of all *p*_*i*_. 2) multiplying the classify probability from 1) with the detection probability from detection step as the final predicted result of the CAD system, if the value ≥ *T*_2_, input will be denoted as true alarm; conversely, as false alarm. During the experiments, *T*_1_ and *T*_2_ are set as 0.5.

**Fig 4.**
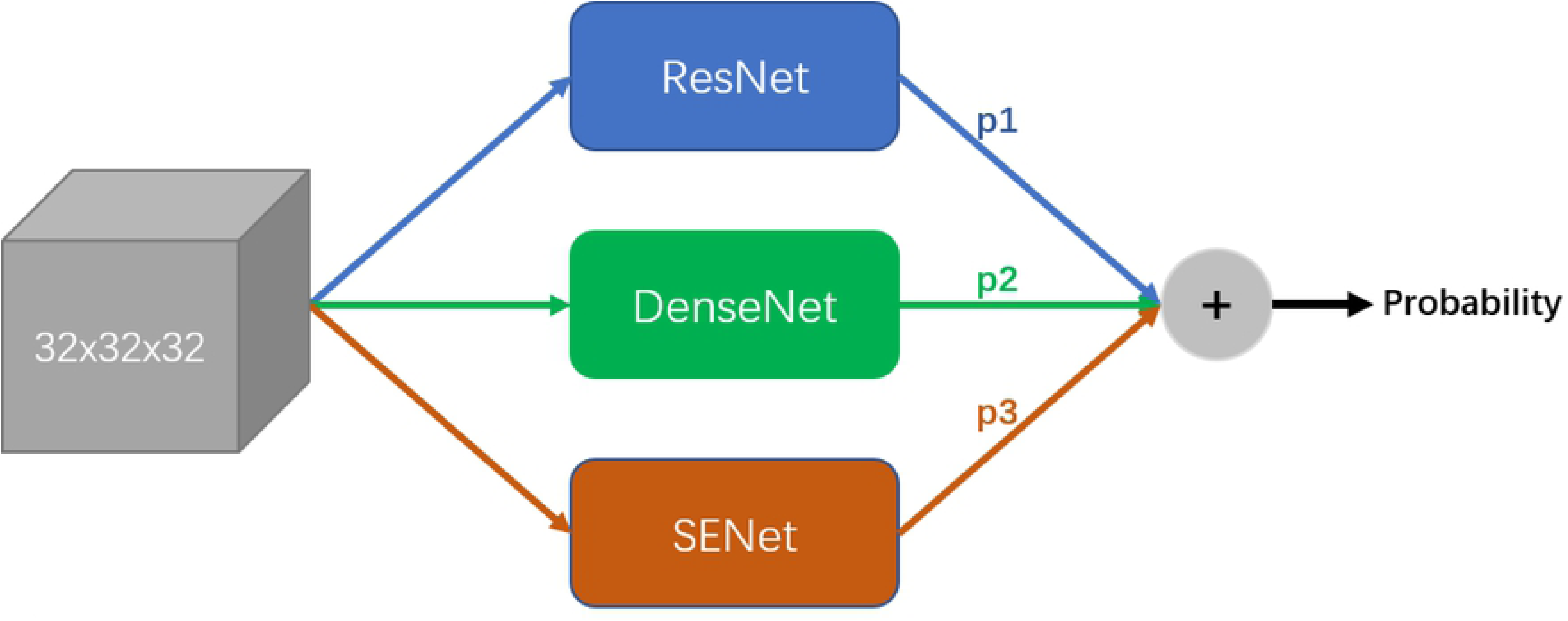
The Architectural Diagram for False Positive Reduction. 1) the input size of the false positive reduction module is 32 × 32 × 32. 2) p1, p2 and p3 are the output probabilities of ResNet, DenseNet and SENet respectively. 3) the final probability is the average value of p1, p2 and p3.

## Experiments

### Dataset

In this paper we employed three lung nodule datasets in all.

- **LUNA16** Dataset [9] comes from LIDC-IDRI [27] which consists of 1018 cases from several institutions. The dataset excluded scans with a slice thickness greater than 2.5 mm. In total, 888 CT scans are included. For each CT scan, nodule candidates are provided and the corresponding class labels (0 for non-nodule and 1 for nodule) are also annotated. 1186 nodules are annotated across 601 patients.
- **LNDb** Dataset [10] contains 294 CT scans collected at the hospital CHUSJ in Porto, Portugal between 2016 and 2018. The dataset excluded scans with a slice thickness greater than 1 mm. All CTs were annotated by at least one radiologist. Finally, the dataset includes 1897 annotations by the 5 radiologists with 1429 corresponding unique findings.
- **ChestCT2019** Dataset [11] is proved by TIANCHI medical AI competition. We get 1397 samples with nodule from the CT scans of 1000 patients. The slice thickness of CTs is less than 2 mm. The biggest difference between ChestCT2019 and the two above datasets is that the slice spacings of CTs in ChestCT are almost all 5 mm.

### Data Preprocessing

The HU values commonly observed in pulmonary nodules in CT scans are used. The Hounsfield scale of tissue density is based on two values: air as −1000 HU and water as 0 HU. Density of other tissues is related to this range, usually from −1000 to +400 HU for pulmonary CT image analysis. Because in pulmonary CT scan exist abundant tissues, in order to decrease the impact of other tissues and avoid losting more information details after we compress raw images into 256-bit grayscale images, we selected [-1350, 150] as the observation range. As the images were collated from different institutions using different CT scanners which results in a wide variation of image parameters within our training dataset, we rescaled the slice thickness of all used CT scans to 1.0 mm.

### Implementation Details

#### Lung Segmentation

The training data of lung cavity segmentation comes from LUNA16. The original label contains five types, background, left lung, right lung, and nodules etc. For lung cavity segmentation, we first merged the label = 3, label = 4 and label = 2 as the lung cavity, label = 1 and label = 5 as the non-lung cavity, and the samples with false label were eliminated. Then, a single CT slice image was selected along the axial direction as input for training. To effectively reducing the over-segmentation rate, when selecting 2D slice images, we set the proportion of slice including and excluding the lung cavity to 7:3. Prevent model overfitting and enhancing its generalization ability, in data augmentation stage we taken some measures like 1) random flipping horizontally or vertically, 2) adding random Gaussian noise, and 3) standardization with the mean of 0 and the variance of 1. During the training, SGD with momentum of 0.9 and initiative learning rate 0.001 is used as optimizer to minimize the our loss function.

#### Pulmonary Nodule Extraction

##### a) 2D NestedUNet for Pulmonary Nodule Extraction

The dataset that we use in this task is from LIDC-IDIR [27], because it contains the particular and relative precise description of pulmonary nodules compared with other datasets, which is helpful to build an accurate segmentation model. For making suitable data as input to network, a large number of patches are randomly cropped from 2D slices according to the center coordinates and sizes of the nodules, whose size is 256 × 256 and must contain complete nodules within the region. To reduce the false positives, we also select the patches which have no nodules for training. In addition, the approaches such as random horizontal and vertical flip, random shift from the crop center, random angle rotation, random color dithering and Gaussian smoothing are used to augment training samples. During the training, SGD with momentum of 0.9 and initiative learning rate 0.001 is used as optimizer to minimize the our BCEDiceLoss (binary cross entropy loss and dice coefficient loss combined with weight).

##### b) 3D RPN for Nodule Extraction

As the 3D RPN network is deep, and has more parameters than 2D RPN, the model is easy to overfit on small dataset. To solve this problem, we use similar methods with above task to enlarge the number of training data. In Section we have discussed the significance of the location of tiny nodule in early pulmonary nodule detection. Hence we increase the sampling frequencies of tiny nodules in the training set. Specifically, the sampling frequencies of nodules larger than 10 mm and 20 mm are expanded 2 and 4 times higher than other nodules, respectively. The sampling frequency of tiny nodules smaller than 3mm also increases 2 times. Some of negative samples have similar appearances with pulmonary nodules, making them difficult to be classified correctly. Hard example mining is a common technique to solve this problem.

#### False Positive Reduction

As for training data of the proposed networks, we firstly rescale each CT in [0, 255] using the specified windowing values, and then normalize it in [0, 1]. After that, for each selected candidate, we use the center of candidate as the centroid and then crop a 64 × 64 × 64 cube. In the training data, there is an imbalance problem on the number of positive and negative samples. To solve the problem, we use the following strategies to augment data:

- Random crop: for each 64 × 64 × 64 cube, we crop smaller cubes in the size of 32 × 32 × 32 from it.
- Rotation: for each 32 × 32 × 32 cube, we rotate it by 90°, 180°, 270° from three orthogonal dimensions (coronal, sagittal and axial position), thus finally augmenting 9 times for each chosen candidate.
- Multiple windowings: for each candidate, we use different windowing values (−1350 to 150 HU, 1200 to 600 HU and −1000 to 400 HU), thus augmenting 3 times.

The training procedure of three classification networks are almost identical. We use SGD with the momentum of 0.9 as optimizer to minimize the loss function and update the model parameters. The initial learning rate is 0.001 and decay is 0.9. The batch size is set to be 128. Due to the imbalance problem of positive and negative samples, we finally choose focal loss [26] as loss function. And the experiments proved that the effect of focal loss is better than cross entropy loss.

## Results

Lung segmentation is very important for analyzing lung related diseases. In this experiment, we compare the performance of our FCNN model with the current methods on dataset [28], including the traditional benchmark method [4], U-Net model [12], R2U-Net [29], and LGAN [30]. 2D Dice score, F1 score and sensitivity were calculated with same settings for each method. The performance comparison is shown in Table 1. Compared with the state of the art methods, our FCNN model has the highest score, with an average F1 score of 0.9836, an accuracy of 0.9965, and an average Dice score of 0.9864. Although the traditional method had the highest specificity, it requires a series of thresholds, morphological manipulations, and composition analysis. Compared with this method, our method provides an end-to-end solution, which takes an average of 3.4 seconds per scan.

**Table 1.** Comparison with Different Pulmonary Nodule Detection Methods on Lung Database [28].

| Methods | F1 score | Sensitivity | Specificity | Accuracy | Dice | Avg. per scan(s) |
| --- | --- | --- | --- | --- | --- | --- |
| F. Liao [4] | 0.9408 | 0.8753 | **0.9994** | 0.9841 | 0.9313 | 63 |
| U-Net [12] | 0.9658 | 0.9696 | 0.9872 | 0.9872 | / | / |
| R2U-Net [29] | 0.9832 | **0.9944** | 0.9832 | 0.9918 | / | / |
| LGAN [30] | / | / | / | / | 0.9850 | / |
| **Our FCNN** | **0.9836** | 0.9834 | 0.9985 | **0.9965** | **0.9864** | 7.7 |

In order to visualize the well performance of our proposed architecture, we compare the predicted results of our FCNN on two different CT slices with the traditional method. Fig 5 shows the segmentation results. The first and second column shows the original input CT slice and the corresponding ground truth, and the other columns display the segmentation results from traditional method and our FCNN respectively. The significant improvement in lung segmentation by using our FCNN model can be observed.

**Fig 5.**
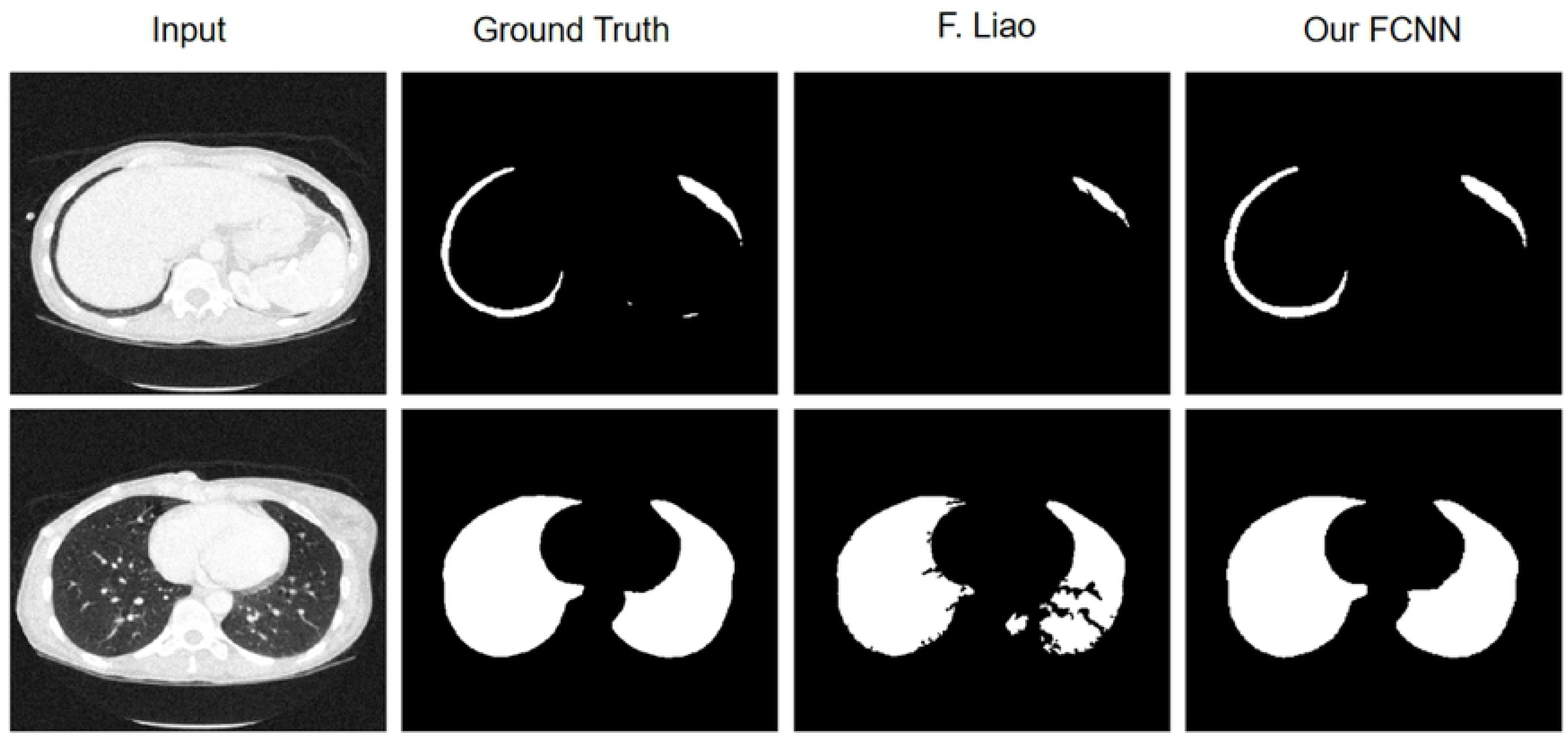
Comparison with Traditional Method from F.Liao [4] and Our FCNN. The first column lists two different scans and their corresponding ground truth in second column. In the third column are the segmentation results by F. Liao and in last column are the results by our method.

We trained the detection model on LUNA16 trainset and evaluated the performance on the validation set of LUNA16. LNDb and ChestCT2019 are also evaluated. In addition, we compared the performance with the DSB and DeepLung models on the three dataset [4, 5]. DSB model from Data Science Bowl 2017 pulmonary nodule detection competition and DeepLung from LUNA16 competition. The average recall comparison of the detection models is shown in Table 2.

**Table 2.** The Average Recall Comparison of Detection Models.

| Method | LUNA16 | LNDb | ChestCT2019 | Ave. per scan(sec) |
| --- | --- | --- | --- | --- |
| DSB | 74.40 | 20.05 | 6.94 | 83 |
| DeepLung | 83.40 | 20.83 | 13.31 | 91 |
| Ours(only 3D-RPN) | 84.54 | 34.77 | 14.05 | **70** |
| Ours | **90.40** | **45.57** | **37.01** | 147 |

From the above table, we can see that our model ensemble method performs better on each data set than the others. The average recall of our model ensemble method on the LUNA16 is 90.4%, 45.57% on LNDB and 37.01% on ChestCT2019. The performance of the only 3D-RPN used method is poor than our complete method, but it takes the shortest time. In [4], the average recall of the DSB method model trained on the Data Science Bowl 2017 competition data set is 0.8562, but it is designed to neglect the very small nodules during training, the LUNA16 dataset is not suitable.

## Conclusion

In this paper, we propose a high-performance pulmonary nodule detection system based on stacking of deep convolution networks. We propose a novel full convolution segmentation network based on U-Net for lung segmentation to solve the efficiency problem of the existing pulmonary nodule detection systems. Furthermore, a 3D-RPN detection network and a 2D-NestedUNet segmentation network is stacked to get the high recall and low false positive rate on pulmonary nodule extraction. Finally, a false positive reduction method based on multi-model ensemble is proposed for the further classification of nodule candidates. Extensive experimental results on public available datasets, LUNA16, LNDb and ChestCT2019, demonstrate the superior performance of our CAD system. We believe that our CAD system will be a very powerful tool for early diagnosis of pulmonary cancer.

## Acknowledgments

The authors thank for the help of Basic Public Research Project of Zhejiang LY20H180013.

